# CDKL5 deficiency results in atypical subregion-specific expression of perineuronal nets during mouse visual critical period

**DOI:** 10.1101/2022.08.31.505900

**Authors:** Brett Addison Emery, Matthew Everett, Logan Reid Dunn, Billy You Bun Lau, Keerthi Krishnan

## Abstract

Perineuronal nets (PNNs) in the primary visual cortex (V1) are specialized extracellular matrix structures that form predominantly on parvalbumin+ GABAergic neurons, marking the closure of visual critical period plasticity. More recently, PNNs are used to characterize deficits in critical period plasticity in mouse models for neurodevelopmental disorders such as Rett syndrome, Fragile X syndrome, and CDKL5 deficiency disorder. Within the mouse V1, studies typically focus on the expression and function of PNNs within the binocular zone, though PNNs are expressed in other subregions of the V1. The expression and role of these PNNs in other subregions are unknown. Here, we performed a systematic whole V1 characterization of PNN expression using *Wisteria floribunda* agglutinin (WFA) staining, with hemisphere-, subregion-, and anatomical axes-specificity, using a null male mouse model for CDKL5 deficiency disorder during the visual critical period. Patients with CDKL5 deficiency disorder often exhibit cerebral cortical visual impairment, though the underlying mechanisms are unclear. Compared to wild-type controls, *Cdkl5*-null males show increased WFA expression at both P15 and P30, with nuanced differences in the subregions, suggesting precocious increase in PNN expression in the *Cdkl5*-null V1. In both genotypes, the binocular zone has significantly higher density of PNNs at both ages, compared to the monocular zone and the rostral V1. These results lay the groundwork to probe the roles for PNNs beyond the binocular zone and cumulatively suggest that, during visual critical period, subregion-specific variations in PNN expression may lead to functional consequences within the *Cdkl5*-null cortex.

## INTRODUCTION

Cortical circuits are shaped by experience during sensitive time windows of early postnatal life. The postnatal maturation of cortical interneurons, specifically parvalbumin+ GABAergic neurons, is critical for controlling the timing of critical period plasticity (Hensch, 2005; Hensch et al., 1998; Huang et al., 1999; Wang et al., 2010). This experience-dependent process is concomitant with the expression of specialized extracellular matrix structures called perineuronal nets (PNNs) on parvalbumin+ GABAergic neurons (Begum and Sng, 2017; Celio, 1993; Gundelfinger et al., 2010; Hartig et al., 1992; Hou et al., 2017; Kosaka and Heizmann, 1989; Krishnan et al., 2015; Miyata and Kitagawa, 2017; Nakagawa et al., 1986; Orlando et al., 2012; Sigal et al., 2019; Sorg et al., 2016; Ueno et al., 2018; Ye and Miao, 2013). PNNs are mainly composed of lecticans, chondroitin sulfate proteoglycans, hyaluronan glycosaminoglycan chains, and other secreted extracellular matrix glycoproteins (Bignami et al., 1992; Carulli et al., 2007; Kwok et al., 2010; Miyata and Kitagawa, 2017). *Wisteria floribunda* agglutinin (WFA) is commonly used as a marker to detect PNNs in the cortex and other brain regions (Brückner et al., 1996; Hartig et al., 1992). WFA specifically binds to N-acetyl galactosamine found on most chondroitin sulfate side chains. Mature PNNs are thought to modulate experience-dependent plasticity (Levelt and Hübener, 2012; Sorg, et al., 2016; Takesian and Hensch, 2013; Wingert and Sorg, 2021), as their increase in developing binocular zone of the primary visual cortex correlates with the termination of the critical period, and PNN removal in adult primary visual cortex restores plasticity, as measured by ocular dominance plasticity assays (Bavelier et al., 2010; Pizzorusso et al., 2006, 2002). In addition to the binocular zone of the primary visual cortex, PNNs are also expressed in other subregions of the primary visual cortex (**Fig. 1**). Currently, their expression during the critical period is unknown.

**Figure 1:**
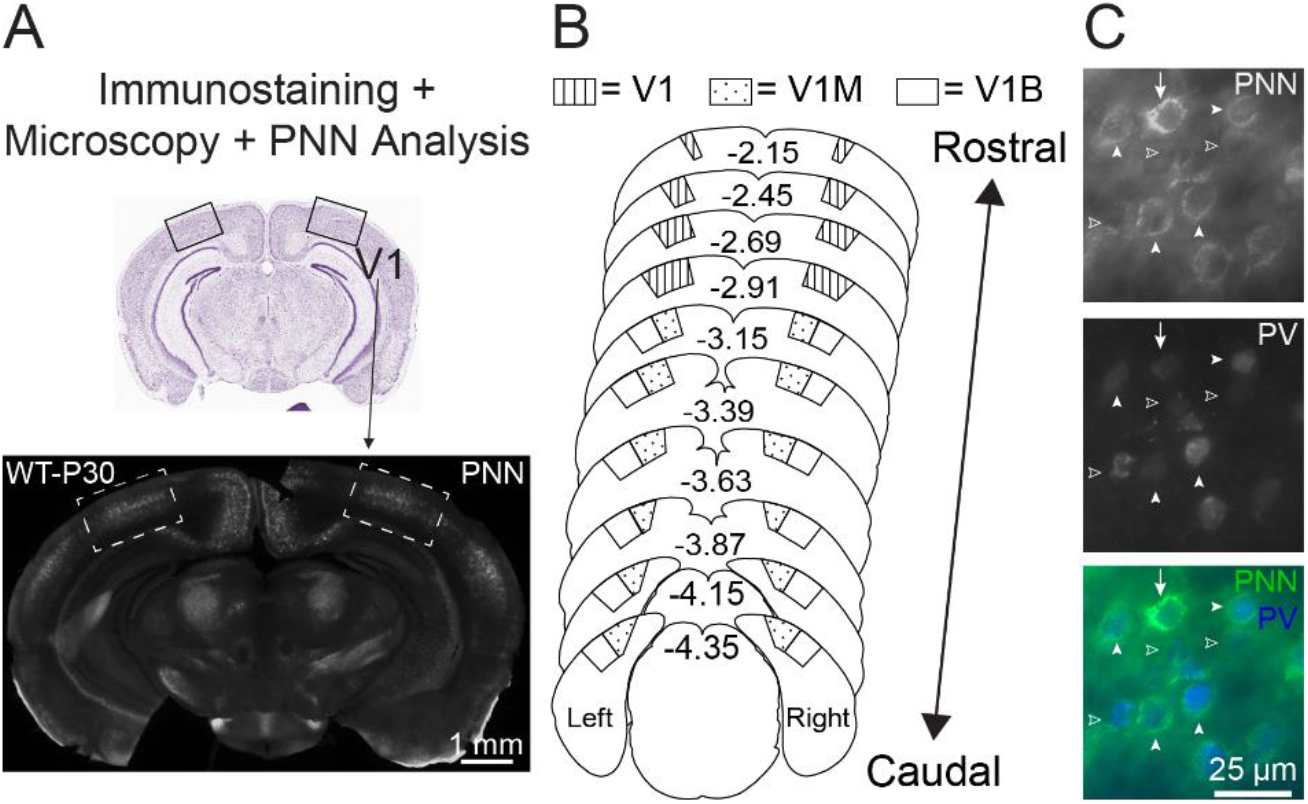
PNN expression analysis across the entire primary visual cortex. **(A)** Top – Nissl-stained coronal mouse brain section, highlighting rostral primary visual cortex (V1; boxed). Bottom: representative coronal wild-type (WT) section immunostained for wisteria floribunda agglutinin (WFA) to identify PNNs (white) in V1 (boxed). In this study, we analyzed postnatal day (P) 15 and P30 brains of WT and CDKL5-null (not shown here). **(B)** Schema of coronal mouse brain sections, depicting rostral to caudal visual cortical subregions analyzed for this study (V1 = rostral V1, V1M = monocular V1, V1B = binocular V1). Values indicate Bregma coordinates according to Paxinos and Franklin, 2013. **(C)** Representative epifluorescent images showing PNNs structures (top; bottom-green) colocalize with parvalbumin+ neurons (PV, middle; bottom-blue) in V1 of P30 WT. Arrow indicates high-intensity PNN. Arrowheads indicate lower intensity PNNs. Open arrowheads indicate diffuse immature WFA signal. All types of PNNs surround PV neurons.

PNN expression is also used as a marker to detect alterations in sensory processing and/or changes in inhibitory networks in mouse models for neurodevelopmental disorders such as Rett syndrome, Fragile X syndrome, and CDKL5 deficiency disorder (Carstens et al., 2021; Donato et al., 2013; Krishnan et al., 2017, 2015; Lau et al., 2020a, 2020b; Patrizi et al., 2020; Pizzo et al., 2016; Reinhard et al., 2019; Wen et al., 2018). CDKL5 deficiency disorder is characterized by seizures, intellectual disability, motor deficits, and social impairments (Brock et al., 2021; Demarest et al., 2019; Siri et al., 2021; Terzic et al., 2021; Zhu and Xiong, 2019). As CDKL5 is on the X chromosome, majority of the children with CDKL5 disorder are girls, though boys are also affected (Bahi-Buisson et al., 2012; Bahi-Buisson and Bienvenu, 2012; Fehr et al., 2016, 2013; Guerrini and Parrini, 2012; Kalscheuer et al., 2003; Weaving et al., 2004). Patients often exhibit cerebral visual impairment (Brock et al., 2021; Mari et al., 2005; Quintiliani et al., 2021), though the underlying mechanisms are unclear. Mouse models of the CDKL5 deficiency disorder recapitulate some of these symptoms, with persistent reduction in response amplitude, reduced visual acuity, and defective contrast function (Mazziotti et al., 2017). Interestingly, these deficits in visually evoked responses are observed in young adult mice (~P60-80) and at the peak of visual critical period plasticity around postnatal day 28 (~P28), but not before (~P25). These results correlate with increased PNN expression within the binocular zone of the *Cdkl5*-null males at ~P35 (Pizzo et al., 2016), suggesting that the timing of critical period plasticity and the proper wiring of visual cortex circuitry are likely affected in rodent models for CDKL5 deficiency disorder.

Though the role of cortical PNNs was established by pivotal studies in the mouse primary visual cortex, all of these studies cited above, performed immunostaining with representative brain sections and mainly focused on the binocular zone (V1B). A systematic characterization of PNNs in the whole primary visual cortex during critical period is crucial to determine the role of PNNs in establishing and/or maintaining visual cortex function. This characterization is also critical to interpret other functional analysis such as visually evoked potentials, which are not typically restricted only to the binocular zone and are relevant as tools for atypical cortical activity in neurodevelopmental disorders (Demarest et al., 2019; LeBlanc et al., 2015; Mazziotti et al., 2017; Wang et al., 2012). Thus, we performed a systematic characterization of PNN expression and density across the developing visual cortex of wild-type (WT) and *Cdkl5*-null (null) mice during visual critical period. In both genotypes, the binocular zone has significantly higher density of PNNs at both ages, compared to the monocular zone and the rostral V1. Manual counting of PNN structures show significantly higher density of high-intensity PNNs in V1B, and a mild reduction in all-intensity PNN density in the V1M of null cortices, compared to wild-type controls at P30. Furthermore, pixel intensity analysis show that *Cdkl5*-null males show increased WFA expression at both P15 and P30, with nuanced differences in the subregions and hemispheres. Together, these results show a precocious increase in PNN expression in the *Cdkl5*-null primary visual cortex during the critical period, suggesting that appropriate expression of CDKL5 is critical for neuronal plasticity during early development, setting the stage for studying PNN expression and function over age, with subregion specificity in the mouse primary visual cortex.

## MATERIALS & METHODS

### Animals

We used the following mouse strains from The Jackson Laboratory: *Cdkl5* hemizygous (B6.129 (FVB) - Cdkl5tm1.1Joez/J; null) (Wang et al., 2012) and wild type (WT) (C57BL/6J). Young male mice (P15, n = 3 animals per genotype; P30, n=5 per genotype) were maintained on a 12-hour light-dark cycle (lights turned on at 07:00 am) and received food ad libitum. Procedures were conducted in accordance with the National Institutes of Health’s Guide for the Care and Use of Laboratory Animals. Protocols were approved by the University of Tennessee in Knoxville’s Institutional Animal Care and Use Committee.

### Immunohistochemistry

WT and null male mice were anesthetized with isoflurane and perfused with 1X phosphate buffer saline (PBS) solution followed by 4% paraformaldehyde (PFA) dissolved in PBS. Brains were extracted and post-fixed in PFA overnight at 4ºC. Prior to sectioning, brains were treated with 30% sucrose in PBS overnight at room temperature. A cut was made on the left ventral-medial brain hemisphere to denote orientation during subsequent tissue processing and analysis. A freezing microtome was used to cut coronal brain sections at 70.0 µm. Sectioning and subsequent histological and imaging steps were done in cohorts, consisting of one WT and one null, to minimize technical variations. Immunohistochemistry was performed as previously described (Krishnan et al., 2015). Briefly, free-floating brain sections were blocked in 10% normal goat serum (NGS) and 0.5% Triton-X in PBS for 3 hours. Then, sections were incubated overnight with biotin conjugated Lectin from *Wisteria floribunda* agglutinin (L1516, Sigma-Aldrich) (1:500) in a 5% NGS and 0.25% Triton-X in PBS. Some tissues were also co-incubated with PV antibodies (mouse, 1:1,000) (P3088, Sigma-Aldrich). The next day, sections were incubated for 4 hours with Streptavidin AlexaFluor-488 (S32354) and anti-mouse CY5 (A10524) (ThermoFisher Scientific; 1:1,000) where appropriate in a 5% NGS and 0.25% Triton-X in PBS. Sections were counterstained with the DAPI (1:1000) for five minutes and mounted on slides in Fluoromount-G for imaging.

### Image Acquisition

Single-plane PNN images of the entire visual cortex from both hemispheres of each brain section were acquired using an epifluorescence microscope (Keyence BZ-X710; Keyence Corp.) equipped with a 10X objective and motorized stage. Imaging settings were determined for each cohort of animals based on the null to minimize overexposure, as preliminary observations suggested brighter PNN expression in null tissues. One cohort consisted of one null and one age-matched wild-type control. Exposure settings for each cohort were determined as previously described (Lau et al., 2020b). Briefly, we identified the exposure time that gives rise to the fewest number of saturated pixels within frame for each tissue. Then, we decreased the exposure time by 1 unit. We completed this process for all null tissues within the cohort and averaged them to get a final exposure time and used this averaged time for final image acquisition of both wild-type and null tissues within the cohort. Images were stitched using the Keyence BZ-X Analyzer software.

### Quantitative Image Analysis

PNN analysis for each hemisphere was performed in ImageJ. We analyzed 10–15 coronal brain sections (both hemispheres) per brain. Rostral V1 (V1), binocular zone (V1B), and monocular zone (V1M) of each image were identified and outlined by overlaying maps from the Paxinos and Franklin’s mouse brain atlas (Paxinos and Franklin, 2013). High-intensity PNNs, representing the most mature PNNs, were counted as previously described (Lau et al., 2020b). Briefly, the contrast setting from ImageJ was maximized to remove weaker signals from the images. The remaining signals were manually quantified as a mature PNN if it retained 80% of its original shape (before contrast adjustment). To determine 80% for each high-intensity PNN, we used the maximized contrast image and the segmented line tool to measure the perimeter of the partial PNN. In the original image, the tips of the segmented line were then connected with a new segmented line to fully encompass the whole PNN. Finally, percentage was calculated by dividing the perimeter value obtained from the partial PNN by the total perimeter (sum of the 2 segmented lines). Detailed protocol and video for mapping and high-intensity PNN analysis can be found in Protocols.IO (https://www.dx.doi.org/10.17504/protocols.io.bcf8itrw).

All PNNs (including high-intensity PNNs) were manually quantified from the ‘original’ image (no contrast nor brightness adjustment); all PNNs had 100% complete borders. To determine left-right asymmetry of PNN expression (**Fig. 5**), for each brain, we averaged the PNN densities (all-PNN or high-intensity PNN) from the left hemisphere and averaged the PNN densities from the right hemisphere. Then, we divided the averaged PNN density value of the left hemisphere by the averaged PNN density value of the right hemisphere to determine the level of PNN asymmetry. For histogram and mean intensity analyses (**Fig. 6-7**), individual intensity values within each subregion region-of-interest (ROI) were acquired from ImageJ by selecting “Analyze” → “Tools” → “Save XY Coordinates” and analyzed in GraphPad Prism. In **Figures 6 and 7**, we illustrate the different histogram ranges in the PNN image after conversion to grey scale. The ranges of intensity were set under the Threshold option in ImageJ. The conversion of intensity ranges between weighted greyscale images and the green signal histogram was performed using the following equation: grey = 0.587green.

### Statistical Analysis

For graphical figures except Figure 5, the analyses were performed using combined hemisphere data. For Figure 5, analyses were performed on parsed left and right hemisphere data. We used Mann-Whitney tests for pair-wise comparison between wild-type and null brains (**Fig. 3C-E, 4, 6, and 7**) and Kruskal-Wallis followed by Uncorrected Dunn’s tests for comparisons between more than 2 groups (**Fig. 2, 3F-H and 5**). The threshold for significance was set at 0.05. Where appropriate, the numbers of animals and images are indicated within the figure legends. Statistical analyses were performed in GraphPad Prism and figures were generated in Adobe Illustrator.

**Figure 2:**
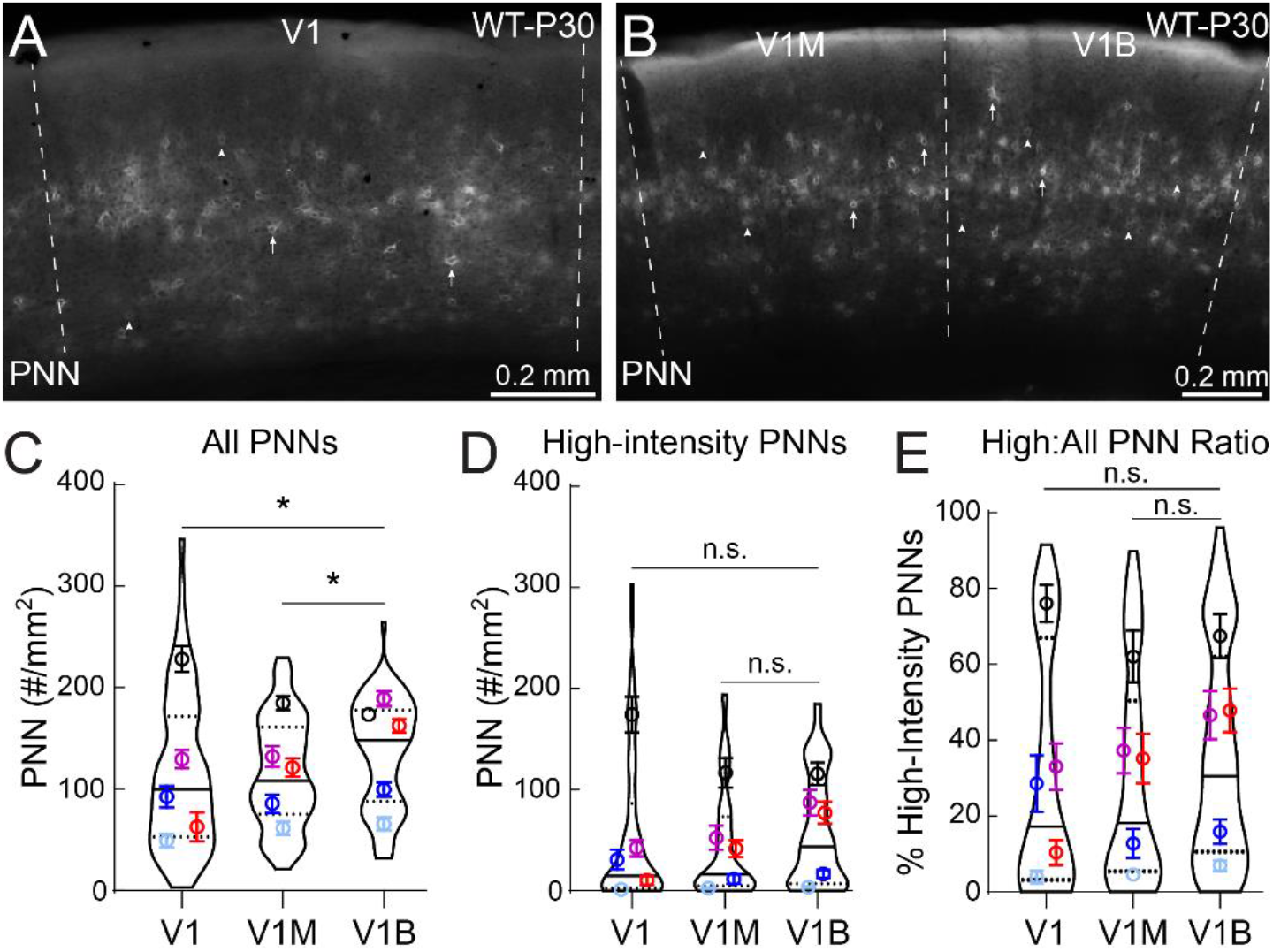
Binocular zone of V1 exhibits higher PNN density than other subregions at P30. **(A-B)** Representative epifluorescent images of PNN expression in rostral V1 (V1, A), monocular (V1M) and binocular (V1B) zones (B). Arrowheads and arrows indicate lower intensity and high-intensity PNN structures, respectively. **(C)** V1B exhibited significantly higher all-PNN density, compared to V1M and V1 (*Kruskal-Wallis followed by Dunn’s test: *p^V1B vs V1M^ = 0*.*037, *pV1B vs V1 = 0*.*013, pV1 vs V1M = 0*.*62)*. **(D)** No significant (n.s.) differences were observed in high-intensity PNNs between the subregions (*Kruskal-Wallis followed by Dunn’s test: p^V1B vs V1M^ = 0*.*075, pV1B vs V1 = 0*.*094, pV1 vs V1M = 0*.*98)*. **(E)** Statistical analysis of high-intensity density to all-PNN density ratio revealed no significant (n.s.) differences between the subregions (*Kruskal-Wallis followed by Dunn’s test: p^V1B vs V1M^ = 0*.*12, pV1B vs V1 = 0*.*16, pV1 vs V1M = 0*.*93)*. For C-E, V1 (64 images), V1M (76 images), V1B (76 images), 5 animals per subregion. Super plots show median (solid line), 25th and 75th quartiles (dash lines) with maximum and minimum, width of violins represents frequency of data points in each region. Each colored circle + lines represent mean ± S.E.M. for a cohort of animals.

**Figure 3:**
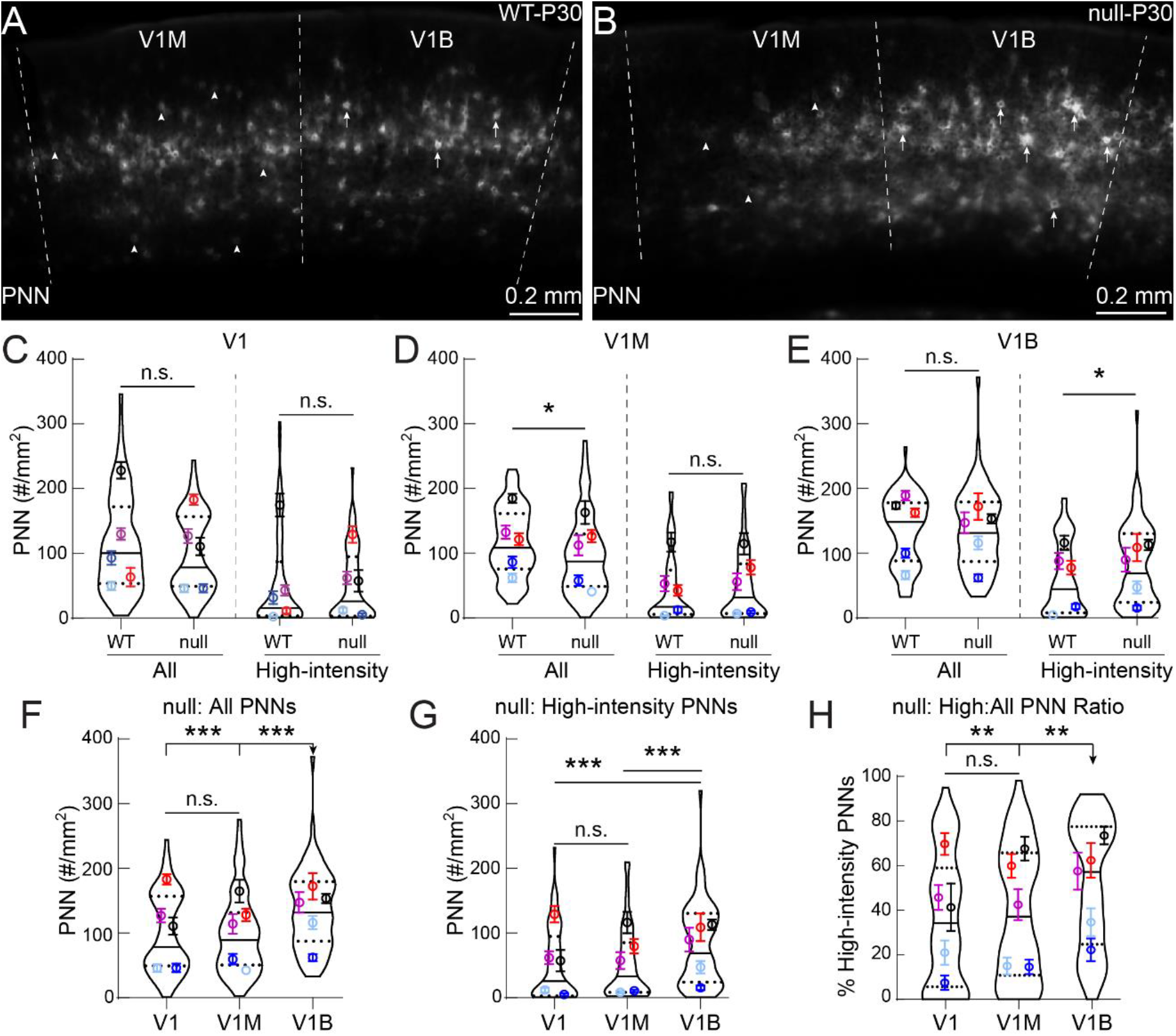
CDKL5-null male mice exhibit atypical PNN expression patterns in subregions of the primary visual cortex at P30. **(A-B)** Representative epifluorescent images of PNN expression at P30, showing CDKL5-null (null) (B) have fewer PNNs in V1M and more high-intensity PNNs in V1B than WT (A). Arrowheads and arrows indicate lower intensity and high-intensity PNNs, respectively. **(C)** In rostral V1 (V1), PNN densities (All- and High-intensity PNNs) were not significantly different between WT and null (WT: n = 64 images; null: n = 62 images; 5 animals per subregion; *Mann-Whitney test: All-PNNs - p = 0*.*26, High-intensity - p = 0*.*88*). **(D)** However, in V1M, null showed significantly decreased density for all-PNNs, with no significant change in high-intensity PNN density, compared to WT (*Mann-Whitney test: All-PNNs - *p = 0*.*013, High-intensity - p = 0*.*48*). **(E)** In V1B, null did not show significant change in all-PNN density, but expressed significantly more high-intensity PNN density compared to WT (*Mann-Whitney test: All-PNNs - p = 0*.*63, High-intensity - *p = 0*.*019*). **(F, G)** Null V1B exhibited significantly higher all-PNN density (F) and higher high-intensity PNN density (G), compared to V1M and V1 (*Kruskal-Wallis followed by Dunn’s test: F - ***pV1B vs V1 = 0*.*0005, ***p^V1B vs V1M^ < 0*.*0001, pV1 vs V1M = 0*.*75; G - ***pV1B vs V1 = 0*.*0003, ***p^V1B vs V1M^ = 0*.*0004, pV1 vs V1M = 0*.*83)*. **(H)** Null V1B exhibited a significantly higher high-intensity PNN density to all-PNN density ratio, compared to V1 and V1M (*Kruskal-Wallis followed by Dunn’s test: **pV1B vs V1 = 0*.*001, **p^V1B vs V1M^ = 0*.*005, pV1 vs V1M = 0*.*54)*. For D and E, WT: n = 76 images, null: n = 72 images; 5 animals per subregion. For F-H, V1 (62 images), V1M (73 images), V1B (72 images), 5 animals per subregion. For C-H, n.s. = not significant; super plots: violin plots show median (solid line), 25th and 75th quartiles (dash lines) with maximum and minimum, width of violins represents frequency of data points in each region. Each colored circle + lines represent mean ± S.E.M. for a cohort of animals.

## RESULTS

We performed a systematic characterization of PNN expression across the subregions of the primary visual cortex by collecting most coronal brain sections covering the entire visual cortex (**Fig. 1A, B**). The primary visual cortex encompasses ~2.2 mm of the mouse brain (Bregma coordinates -2.15 mm to -4.35 mm (Paxinos and Franklin, 2013)) and is composed of the rostral V1 (V1), monocular zone (V1M), and binocular zone (V1B) subregions (**Fig. 1B**). Within the cortex, PNNs surround the soma and proximal dendrites of parvalbumin+ GABAergic neurons (Agetsuma et al., 2018; Bartos et al., 2002; Cardin et al., 2009; Celio, 1993; Cho et al., 2020; Faini et al., 2018; Hartig et al., 1992; Karunakaran et al., 2016; Ko et al., 2013; Sohal et al., 2009; Ueno et al., 2018); PNN expression is predominantly in superficial layers of the cortex (Layers II/III, IV), with minimal expression in deeper layers (Layers V, VI). As assessed by *Wisteria floribunda* agglutinin (WFA) staining (Brückner et al., 1996; Hartig et al., 1992), PNN expression around individual neurons can be strong, weak or diffused (**Fig. 1C**, arrows, arrowheads or open arrowheads, respectively).

During the peak of mouse critical period plasticity (P30), WT males show more distinct PNN expression within the V1M and V1B subregions compared to rostral V1 (**Fig. 2A, B**). Through manual quantification of all PNNs across these subregions, we found that V1B showed significantly higher PNN density compared to V1 and V1M (**Fig. 2C**). However, the density of mature, high-intensity PNNs did not differ between subregions (**Fig. 2D**). The ratio of high-intensity PNNs to all PNNs (represented as a percentage) did not vary significantly between subregions (**Fig. 2E**), suggesting that although PNN density is highest in V1B, the majority of PNNs are not of full maturity. Together, these results show that PNN expression is subregion-specific in WT males, with V1B showing significantly greater PNN density than V1M or rostral V1 following the peak of critical period plasticity.

Next, we determined if the male mouse model for CDKL5 deficiency disorder displayed subregion-specific alterations in PNN expression, as was previously shown in the V1B (Pizzo et al., 2016). We found PNN expression in P30 null males appeared to be distinctly different in multiple visual cortex subregions compared to WT controls (**Fig. 3A, B**). Compared to WT controls, all PNN density within V1M was significantly decreased in P30 null males, while no significant difference in high-intensity PNN density was observed between conditions (**Fig. 3D**). In contrast, high-intensity PNN density within V1B was significantly greater in null than WT, with no significant difference in all PNN density (**Fig. 3E**). We observed no significant differences in PNN densities between conditions in rostral V1 (**Fig. 3C**). While comparing within the null genotype, PNN density in V1B was significantly higher compared to V1M and rostral V1, with higher densities of all PNNs, high-intensity PNNs and ratio of high-intensity to all PNN (**Fig. 3F, G, H, respectively)**. These results are in contrast with the WT profile (**Fig. 2C-E)**. Together, these results suggest that subtle, yet significant subregion-specific differences exist in PNN density within the *Cdkl5*-null mouse visual cortex at P30.

We previously reported that the adult WT female primary somatosensory cortex exhibited anatomical axis- and hemisphere-specific changes in high-intensity PNN density (Lau et al., 2020b), which may contribute to functional specialization within cortical circuitry. To determine if the developing visual cortex of male mice also exhibits such anatomical and hemisphere-specific expression patterns during critical period, we plotted PNN densities (all PNNs and high-intensity PNNs) across the rostral-caudal axis (**Fig. 4**). In contrast to the adult primary somatosensory cortical PNN density distribution, we observed no significant differences in either all or high-intensity PNN density within genotype along the rostral-caudal axis. Compared to WT controls, P30 null males showed significant decreases in all PNN density in the more caudal areas of V1M (**Fig. 4B**) and a significantly increased high-intensity PNN density in the rostral V1B (**Fig. 4F**). We did not observe significant differences in all or high-intensity PNN density between WT and null rostral V1 along the rostral-caudal axis (**Fig. 4A, D**). Together, these results point to specific and limited anatomical disruptions in the *Cdkl5*-null primary visual cortex.

**Figure 4:**
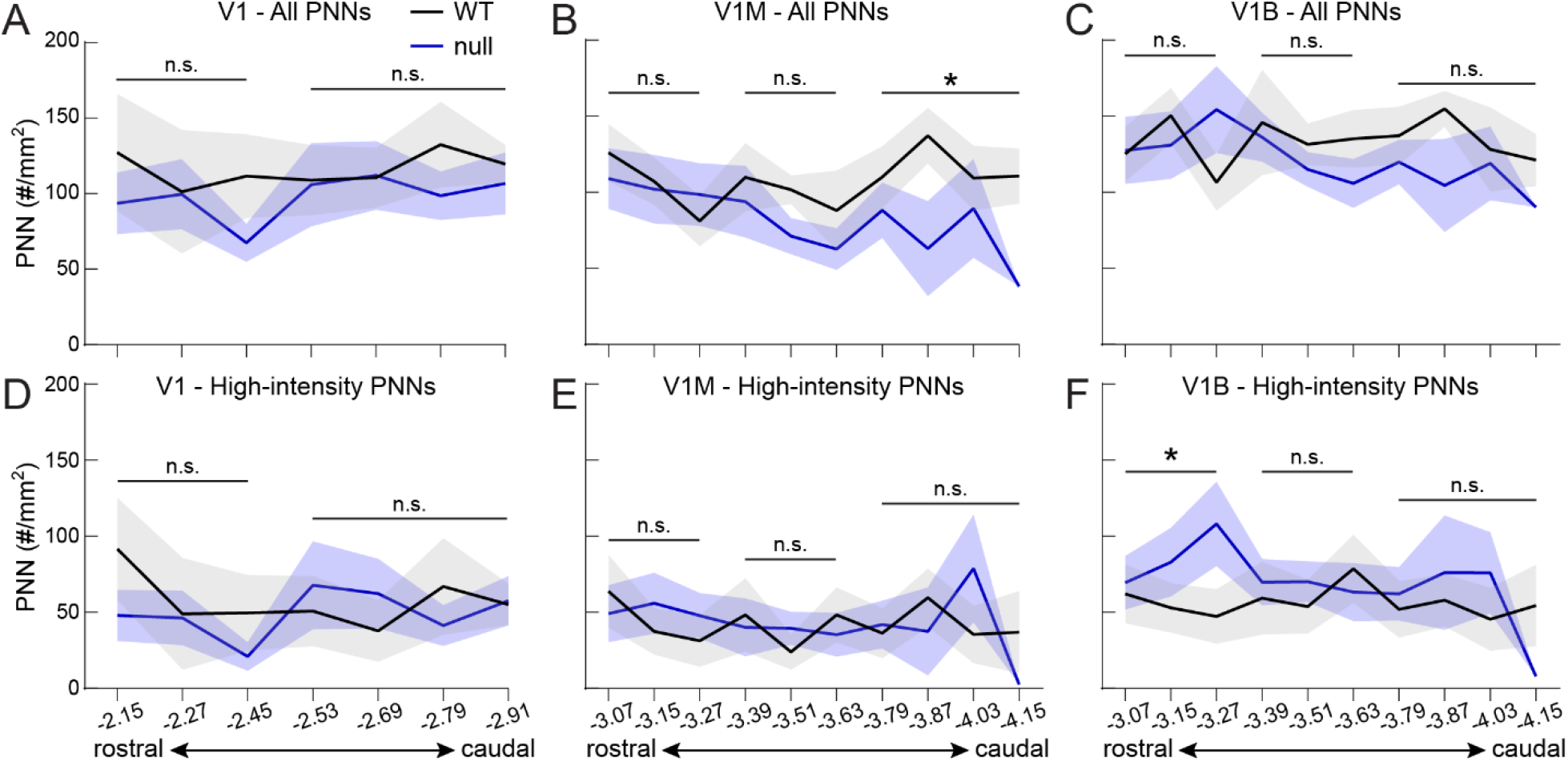
CDKL5-null male mice exhibit altered PNN expression across rostral-caudal axis in specific subregions of primary visual cortex at P30. **(A-F)** Distribution of all-PNN density (A-C) and high-intensity PNN density (D-F) in rostral V1 (A, D), V1M (B, E) and V1B (C, F) of WT (black) and null (blue) (solid lines) across rostral-caudal axis. Null exhibited a significant reduction in all-PNN density in more caudal regions of V1M (B) and a significant increase in high-intensity PNN density in rostral V1B (F), compared to WT. N = 1-13 images per Bregma coordinate, 5 animals per subregion. Sets of combined Bregma coordinates (as indicated by horizontal bars) were used for statistical analysis. Coordinates are based on Paxinos and Franklin, 2013. Solid lines + shades represent mean ± S.E.M. *Mann-Whitney test within each subregion between WT and null*: V1M-All-PNNs: *\*p^WT vs null^ = 0*.*016*, V1B-High-intensity PNNs: *\*p^WT vs null^ = 0*.*032, n*.*s. = no significance*. See Table 1 for full statistical results.

Next, we investigated if there were any hemisphere-specific changes in PNN expression. Plotting PNN density of individual sections between the left and right hemispheres, we observed no significant difference in all PNN (**Fig. 5A, D, G**) or high-intensity PNN (**Fig. 5B, E, H**) density within genotype across all visual cortex subregions. Analyzing left-right asymmetry within individual brains showed remarkable consistency in V1B (**Fig. 5I**), while V1 and V1M showed minor individual variations (**Fig. 5C, F**, red and light blue circles), indicative of similar PNN densities between hemispheres in both grouped and individual analyses. One statistically significant difference was observed between genotypes in the left hemisphere of V1M: all PNNs (**Fig. 5D**), though the overall distribution remains similar. These findings cumulatively suggest that (*1*) hemisphere laterality has not been established within the primary visual cortex during the peak of critical period plasticity, and (*2*) this process is not dysregulated at P30 within the male *Cdkl5*-null visual cortex.

**Figure 5:**
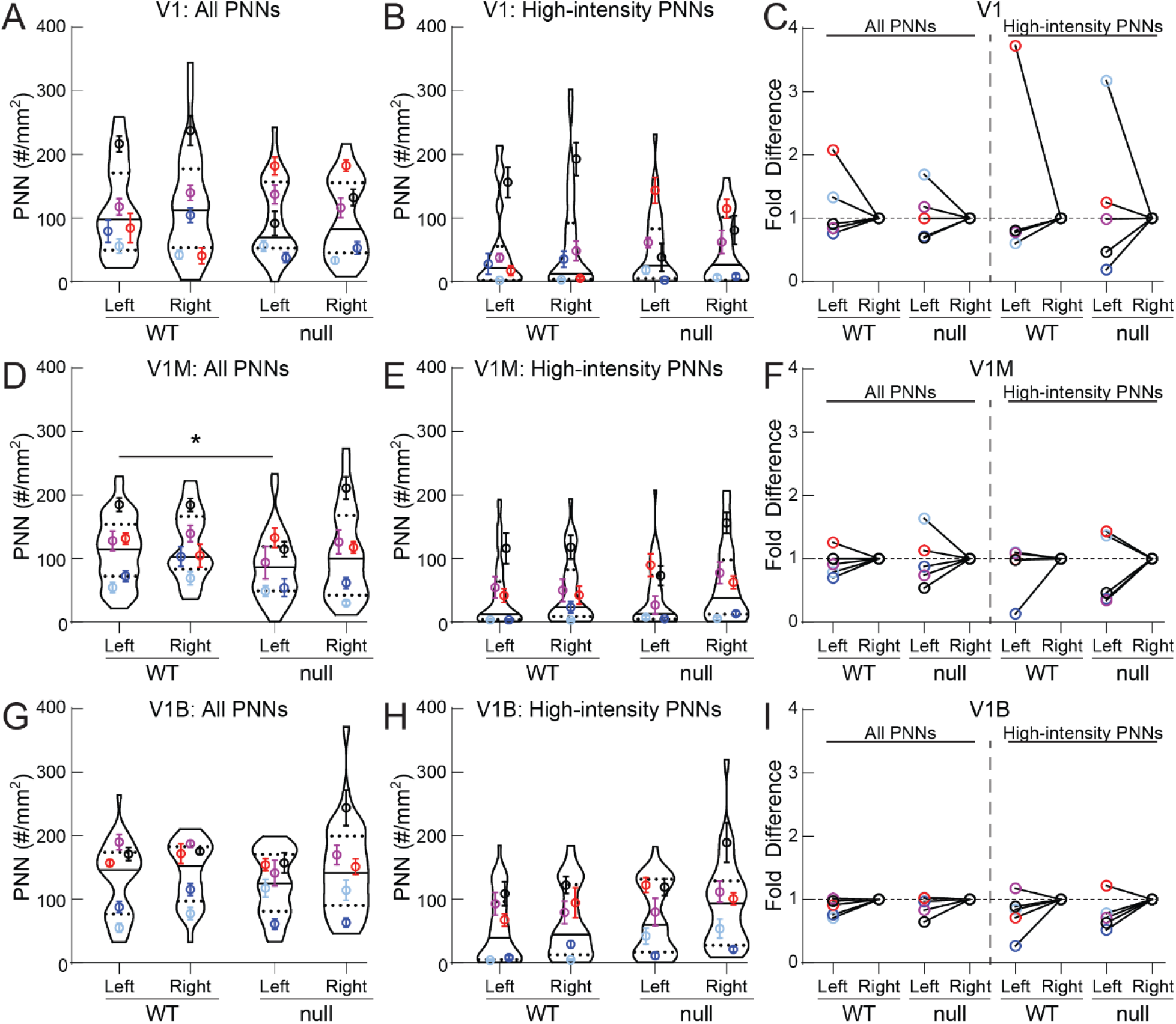
Individual brains exhibit similar PNN density between hemispheres in all primary visual cortical subregions at P30. On average and within genotype, WT and null exhibited similar all-PNN density **(A, D, G)** and high-intensity PNN density **(B, E, H)** between left and right hemispheres of rostral V1 (A, B), V1M (D, E) and V1B (G, H). Overall, there were no hemisphere-specific changes between WT and null visual cortex, except for V1M, where null showed a significant reduction in all-PNN density in the left hemisphere (D) (*Kruskal-Wallis followed by Dunn’s test:* V1M: \**pWT-left vs null-left = 0*.*035*). **(C, F, I)** Individual brains displayed natural variations in asymmetry in both all-PNN and high-intensity PNN densities in rostral V1 (C), and minimally in V1M (F) and V1B (I). For each brain, PNN density from the left hemisphere was normalized to its right hemisphere. (A-B, D-E, G-H) Super plots: violin plots showing median (solid line), 25th and 75th quartiles (dash lines) with maximum and minimum, width of violins represents frequency of data points in each region. Each colored circle + lines represent mean ± S.E.M. for a cohort of animals. N = 31-42 images, 5 animals per group. See Table 2 for full statistical results.

**Table 1:** Statistical p values for Figure 4

|  | V1 |  | V1M |  |  | V1B |  |  |
| --- | --- | --- | --- | --- | --- | --- | --- | --- |
| Bregma values | -2.15 to -2.45 | -2.53 to -2.91 | -3.07 to -3.27 | -3.39 to -3.63 | -3.79 to -4.15 | -3.07 to -3.27 | -3.39 to -3.63 | -3.79 to -4.15 |
| All PNN | 0.58 | 0.36 | 0.85 | 0.069 | **0.016\*\*** | 0.68 | 0.18 | 0.14 |
| High-Intensity | 0.84 | 0.68 | 0.35 | 0.88 | 0.92 | **0.032\*** | 0.71 | 0.46 |

**Table 2:** Statistical p values for Figure 5

|  | Fig 5A<br>All | Fig 5B<br>High | Fig 5D<br>All | Fig 5E<br>High | Fig 5G<br>All | Fig 5H<br>High |
| --- | --- | --- | --- | --- | --- | --- |
| WT: Left vs Right | 0.96 | 0.94 | 0.60 | 0.35 | 0.32 | 0.30 |
| null: Left vs Right | 1.00 | 0.93 | 0.28 | 0.090 | 0.20 | 0.20 |
| Left: WT vs null | 0.40 | 0.92 | **0.035\*** | 0.92 | 0.59 | 0.11 |
| Right: WT vs null | 0.43 | 0.91 | 0.16 | 0.39 | 0.85 | 0.09 |

As we observed an increase in the high-intensity PNN density within V1B of P30 null males, we wondered if the etiology of CDKL5 deficiency disorder was similar to that of the *Mecp2*-null visual cortex, with a precocious increase in high-intensity PNN density, indicative of an accelerated critical period (Krishnan et al., 2015). Thus, we analyzed PNN expression at the start of the critical period, in P15 WT and null brains. WFA staining in developing visual cortex is typically weak and diffuse after eye opening (~P12), with some layer IV neurons displaying a classic net-like structure (Krishnan et al., 2015; Pizzorusso et al., 2002; Ye and Miao, 2013). To capture the WFA staining patterns in this immature state, we performed pixel intensity analysis across individual images using ImageJ. In a representative example, the histogram intensity values (**Fig. 6A, B**) corresponded to varying staining patterns across layers, with (*i*.*-ii*.) intensity value of 1-49 representing fiber staining in corpus callosum and deeper layers of the cortex, (*iii*.) 50-79 representing weak yet specific staining in superficial and deep cortical layers, (*iv*.) 80-124 identifying more defined PNN structures in Layer IV as well as background staining in Layer I, and (*v*.) over 125 identifying high-intensity PNNs, mostly in Layer IV. A binned intensity analysis over all P15 images of WT and null sections showed higher mean intensities in null brains, especially over the weak staining in upper layers (**Fig. 6C**, *iii*.), in all three subregions. There was no statistically significant difference in subregion-specific overall mean intensity measurements between P15 WT and null, though the distributions were distinct (**Fig. 6D**). Together, these results indicate an overall precocious increase in PNN expression in different layers and structures, though not of the typically measured mature high-intensity PNNs.

We performed a similar analysis with P30 brain sections, to relate the histogram analysis to manually quantified PNNs (all and high-intensity PNNs). As seen with P15 tissues, the intensities between 80-124 delineated the most well-formed PNNs in Layer IV (**Fig. 7A, B**, *iv*.), with intensities of 125 and above identifying the high-intensity PNNs (**Fig. 7A, B**, *v*.). In the null rostral V1, we detected a significant increase in the fiber staining in the corpus callosum and deeper layers (*i*), with no overall change in mean intensity (**Fig. 7C, D**, top panels). In the V1M, pixel intensity analysis showed the alternating dynamics in intensity groups (*ii)* and *(iii)* between the WT and null, with significant increases in pixel intensity in (*iv)* and (*v)* groups, corresponding to layers I and IV, of the null V1M (**Fig. 7C, D**, middle panels). These results are in contrast with the observed decrease in all-intensity PNN density in the null V1M, from the manual counting of PNN structures (**Fig. 3D**). In the V1B, pixel intensity analysis showed an increase in mean intensity in the null V1B compared to WT controls, recapitulating the manual high-intensity PNN analysis (**Fig. 7C-*v*, D**, bottom panels vs. **Fig. 3E**). Together, these results reiterate the increase in defined PNN structures in the null V1B, and identify nuance changes at different levels of PNN expression in the V1M.

**Figure 6:**
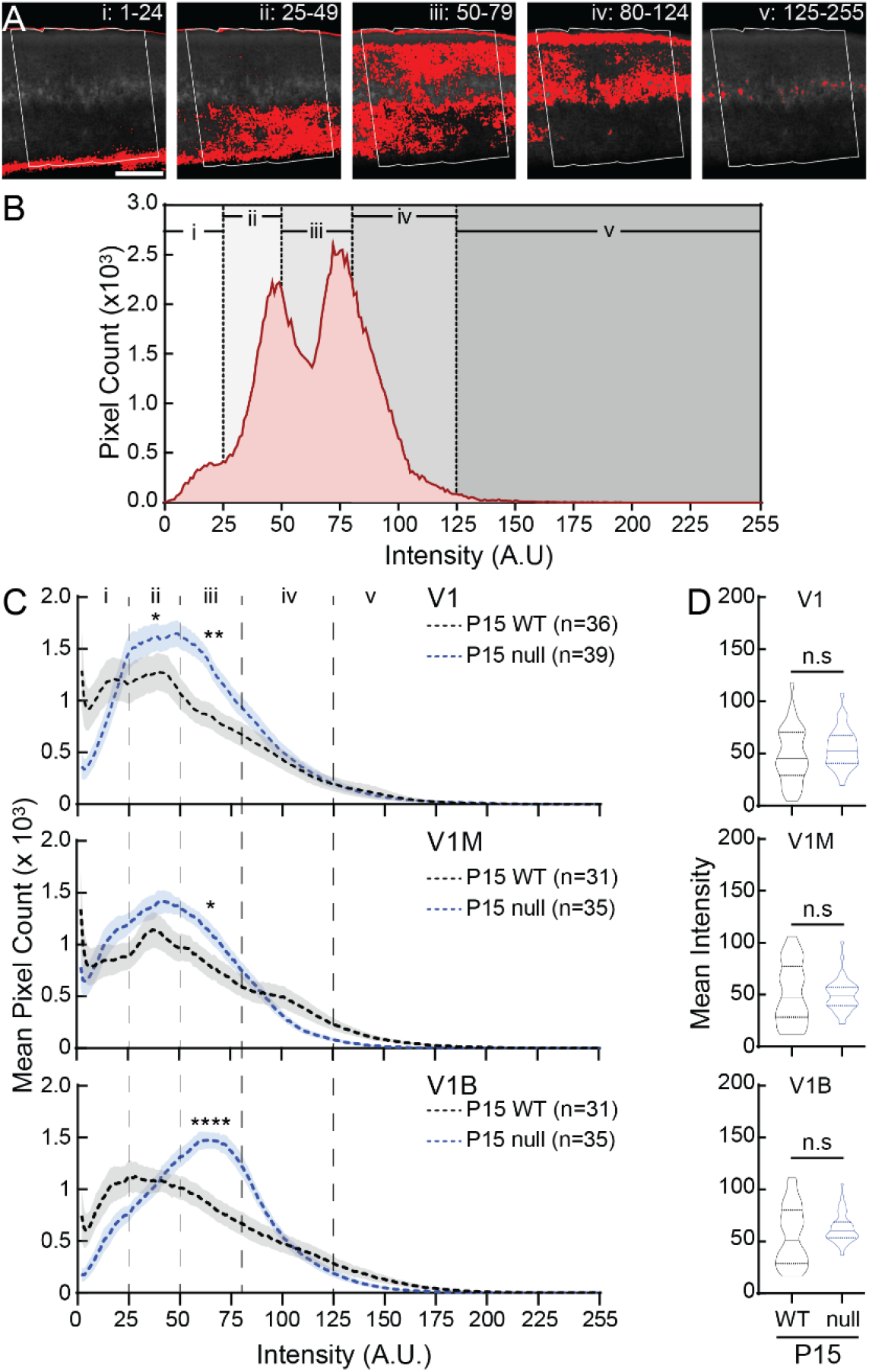
CDKL5-null male mice express increased WFA intensity in all subregions of the primary visual cortex at P15. **(A)** One representative greyscale WFA image in V1B (boxed) is used as an example to show five different ranges of pixel intensity in ImageJ, and their locations in different layers (i-v). Red indicate signals measured at each of the five intensity ranges. **(B)** Histogram showing intensity distribution of all WFA signals from the same image in A. **(C)** Averaged histograms showing WFA intensity distribution in V1 (top), V1M (middle) and V1B (bottom) of WT (black) and null (blue) (dash lines). In V1, null had significantly higher WFA intensities in the ranges of ii and iii compared to WT (*Mann-Whitney test: ranges i, p = 0*.*81; ii, *p = 0*.*019; iii, **p = 0*.*0019; iv, p = 0*.*11; v, p = 0*.*44*). In V1M, null had significantly higher WFA intensities in the range of iii compared to WT (*Mann-Whitney test: ranges i, p = 0*.*32; ii, p = 0*.*17; iii, *p = 0*.*050; iv, p = 0*.*48; v, p = 0*.*94*). In V1B, null had significantly higher WFA intensity in iii compared to WT (*Mann-Whitney test: ranges i, p = 0*.*11; ii, p = 0*.*77; iii, ****p < 0*.*0001; iv, p = 0*.*14; v, p = 0*.*53*). Mean (dash lines) ± S.E.M. (shades) are shown. N values represent numbers of images analyzed for each condition, with 3 animals per group. **(D)** Combined-hemisphere statistical analysis of V1 (top), V1M (middle) and V1B (bottom) revealed no significant (n.s.) differences in mean WFA intensity without the different groupings between WT and null (*Mann-Whitney test: pV1-WT vs V1-null = 0*.*17, pV1M-WT vs V1M-null = 0*.*99, pV1B-WT vs V1B-null = 0*.*19)*. Violin plots showing median (solid line), 25th and 75th quartiles (dash lines) with maximum and minimum, width of violins represents frequency of data points in each region. N values for each group are the same as in C.

**Figure 7:**
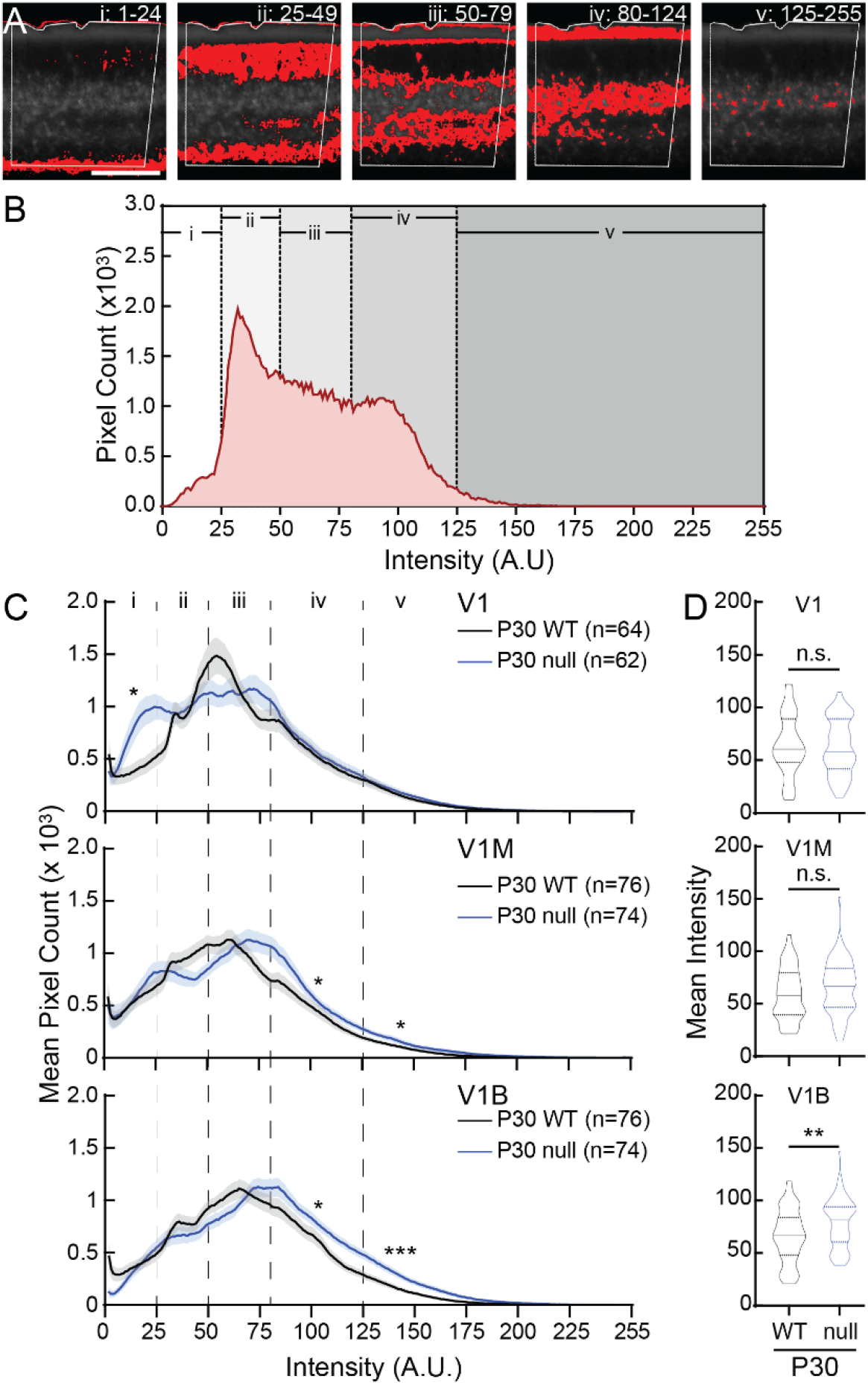
CDKL5-null male mice express increased WFA intensity in a V1 subregion-specific manner at P30. **(A)** One representative greyscale WFA image in V1B (boxed) is used as an example to show five different ranges of pixel intensity in ImageJ and their locations in different layers (i-v). Red indicate signals measured at each of the five intensity ranges. **(B)** Histogram showing intensity distribution of all WFA signals from the same image in A. **(C)** Averaged histograms showing WFA intensity distribution in V1 (top), V1M (middle) and V1B (bottom) of WT (black) and null (blue) at P30 (solid lines). In V1, null had significantly higher WFA intensity in the range of i compared to WT (*Mann-Whitney test: ranges i, *p = 0*.*039; ii, p = 0*.*74; iii, p = 0*.*98; iv, p = 0*.*66; v, p = 0*.*23*). In V1M, null had significantly higher WFA intensity in the ranges of iv and v compared to WT (*Mann-Whitney test: ranges i, p = 0*.*94; ii, p = 0*.*17; iii, p = 0*.*64; *iv, p = 0*.*021; v, *p = 0*.*027*). In V1B, null had significantly higher WFA intensity in the ranges of iv and v compared to WT (*Mann-Whitney test: ranges i, p = 0*.*57; ii, p = 0*.*20; iii, p = 0*.*96; *iv, p = 0*.*011; v, ***p = 0*.*0002*). Mean (solid lines) ± S.E.M. (shades) are shown. N values represent number of images analyzed for each condition, with 5 animals per group. **(D)** Combined-hemisphere statistical analysis revealed no significant (n.s.) differences in mean WFA intensity between WT and null in V1 (top) and V1M (middle). However, null expressed significant higher mean WFA intensity in V1B (bottom) (*Mann-Whitney test: pV1-WT vs V1-null = 0*.*68, pV1M-WT vs V1M-null = 0*.*13, **pV1B-WT vs V1B-null = 0*.*007)*. Violin plots showing median (solid line), 25th and 75th quartiles (dash lines) with maximum and minimum, width of violins represents frequency of data points in each region. N values for each group are the same as in C.

## DISCUSSION

PNNs are widely expressed in the rodent brain; in particular, the different sensory cortical regions display dense PNNs surrounding the soma of neurons. Though their relative functions in individual brain regions and cell types may vary (Balmer, 2016; Bartos et al., 2002; Begum and Sng, 2017; Bernard and Prochiantz, 2016; Carstens et al., 2016; Dauth et al., 2016; de Winter et al., 2016; Deepa et al., 2002; Devienne et al., 2021; Dityatev et al., 2007; Donato et al., 2013; Durand et al., 2012; Frischknecht et al., 2009; Gundelfinger et al., 2010; Hartig et al., 1992; Hou et al., 2017; Kalemaki et al., 2018; Kosaka and Heizmann, 1989; Krishnan et al., 2017, 2015; Lau et al., 2020a; Liu et al., 2021; Nakagawa et al., 1986; Orlando et al., 2012; Pantazopoulos et al., 2020; Pizzorusso et al., 2006, 2002; Sigal et al., 2019; Sugiyama et al., 2009; Suttkus et al., 2012; Tansley et al., 2022; Tewari et al., 2018; Ueno et al., 2018; Vo et al., 2013; Wingert and Sorg, 2021; Ye and Miao, 2013), PNNs generally regulate the function of inhibitory networks during early postnatal development and across life. As PNNs are predominantly found on PV+ GABAergic neurons in the cortex, they are intimately associated with, and contribute to, PV network function underlying cellular and organismal behavior (Agetsuma et al., 2018; Bartos et al., 2002; Cardin et al., 2009; Carstens et al., 2016; Cattaud et al., 2018; Cho et al., 2020; Donato et al., 2013; Faini et al., 2018; Gogolla et al., 2009; Karunakaran et al., 2016; Ko et al., 2013; Krishnan et al., 2017, 2015; Lau et al., 2020a; Miyata et al., 2012; Murthy et al., 2019; Nowicka et al., 2009; Sigal et al., 2019; Sohal et al., 2009; Thompson et al., 2018; Wingert and Sorg, 2021). Determining the function of PNNs with regional and biological contexts requires a thorough characterization of their expression across the whole brain (Lupori et al., 2023; Ueno et al., 2018). Particularly, Ueno et al. (2018) noted an increase in WFA+ PNNs in the mouse V1B from two to six to 12 months of age, with no differences in V1M, from representative section analysis. This study highlights the need for systematic analysis across age and whole-brain regions. Here, we have performed a systematic analysis of PNN expression during the well-accepted critical period of the primary visual cortex (**Fig. 8**). In the early stages of visual cortex critical period development (~P15), WFA expression is generally diffuse in superficial layers, with minimal PNN structures in Layer IV as typically observed across visual cortex subregions (**Fig. 6**). However, by the peak of critical period plasticity at P30, PNN expression is distinct in layers IV/V of V1B, compared to V1M and rostral V1 areas (**Fig. 2, 7**). High expression of PNNs on parvalbumin+ GABAergic neurons in the primary sensory areas could selectively control thalamic excitation onto these neurons, and control of feed-forward thalamocortical sensory inputs into the sensory cortex (Faini et al., 2018; Lupori et al., 2023). Such functional connectivity regulations could have strong impact on the known roles of the binocular zone of the primary visual cortex in ocular dominance plasticity, binocular matching, and depth perception (Antonini et al., 1999; Fagiolini and Hensch, 2000; Krishnan et al., 2015; Pizzorusso et al., 2006, 2002; Wang et al., 2010). However, the functional relevance of PNN expression in V1M and rostral V1 requires further elucidation.

**Figure 8:**
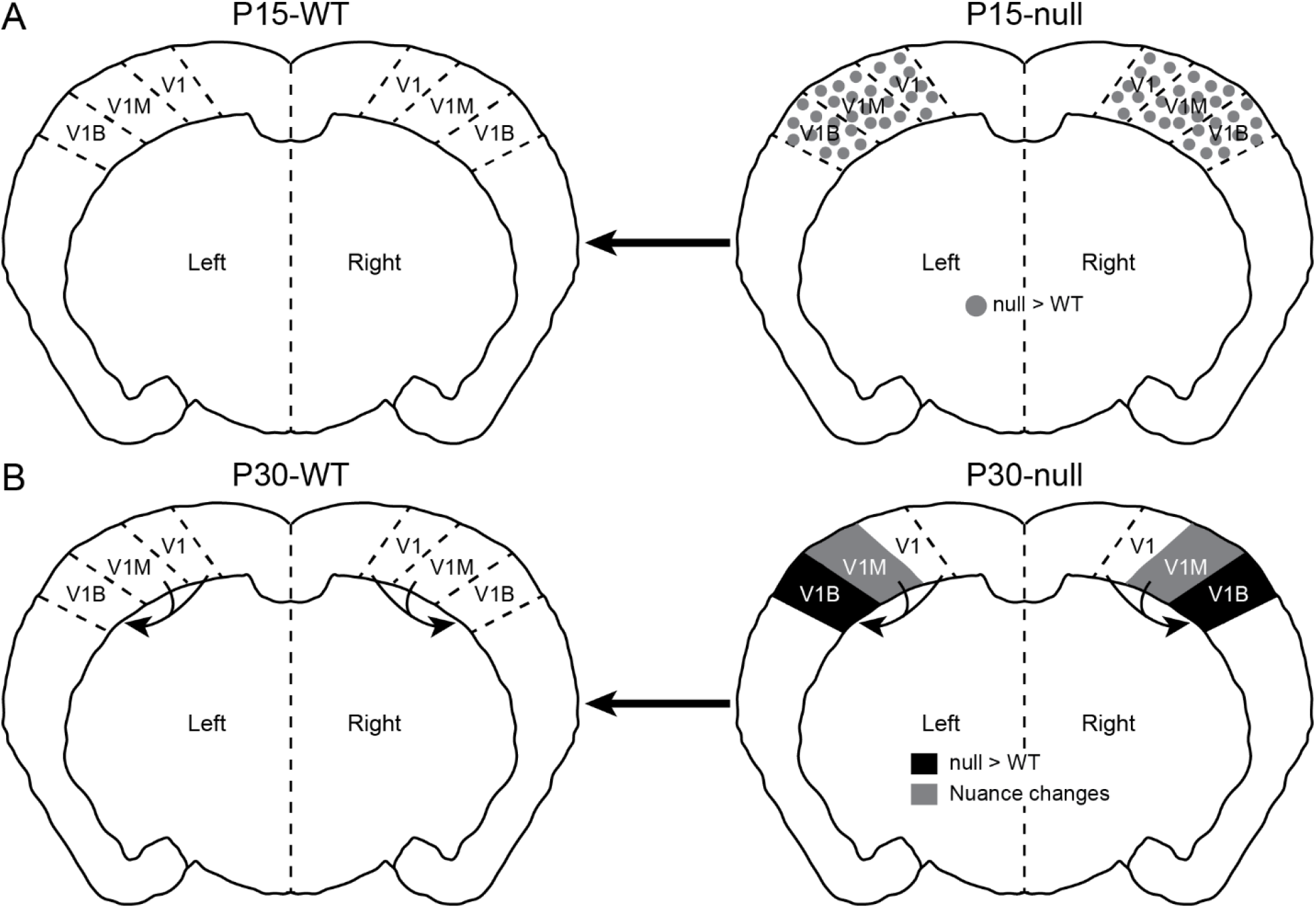
Summary of PNN expression changes between WT and CDKL5-null during critical period in subregions of the primary visual cortex. (A) Before the beginning of critical period at P15, null exhibited increased diffuse WFA expression in V1, V1M and V1B (grey dots), suggesting the beginnings of PNN dysregulation. **(B)** At the peak of critical period, null exhibited increased PNN expression in V1B (black). In V1M, null had nuanced changes in PNN expression compared to WT: null showed a decrease in the density of overall PNN structures, by manual counting, (Fig. 3D and 4B), and a higher intensity of WFA signal (Fig. 7C). Together, these results suggest fluid and compensatory changes in PNN expression in null V1M at these ages. The impact of such changes over time, on synaptic plasticity and visual cortex function, particularly in V1M, needs further work.

Typically, neurons of monocular V1 are thought to encode for receptive field properties including orientation and direction selectivity. However, the role of neuronal activity and/or visual experience in initial formation and maturation of these properties remains controversial (Chapman and Stryker, 1993; Hagihara et al., 2015; Ko et al., 2013; Miller et al., 1999). Furthermore, the role of inhibitory neurons in the establishment and maintenance of receptive field properties in monocular V1 remains to be determined. The emergence of multi-unit recordings and wide-field imaging techniques with natural scenery stimuli in different species could begin to shed light on the role of V1M and, ultimately, on parvalbumin+ interneurons and PNNs across subregions of the primary visual cortex. As proteins involved in forming perineuronal nets are thought to be secreted by different cell types, the systematic coordination, assembly, and maintenance of PNNs during critical periods of development need further characterization. Cortical PNNs predominantly surround parvalbumin+ GABAergic neurons, the latter of which are typically identified through genetic strategies or immunostaining of parvalbumin protein. Though immunostaining is frequently used for quantitative analysis of parvalbumin expression, parvalbumin protein level is regulated in an activity-dependent manner (Berardi et al., 1993; Filice et al., 2016; Krishnan et al., 2015; Patz et al., 2004; Tropea et al., 2006; Ye and Miao, 2013). Thus, caution must be used in interpreting correlations between parvalbumin expression and PNN expression when solely relying on immunostaining techniques.

Neurodevelopmental disorders are synaptic disorders, which are particularly affected in early sensory critical periods when the fully formed brain is being sculpted by experience-driven neural activity. Studies in animal models of neurodevelopmental disorders such as Rett syndrome, Fragile X syndrome, and CDKL5 deficiency disorder have shown precocious or delayed impacts on sensory critical period processes, using both electrophysiological approaches and/or detection of altered PNN expression by immunostaining analysis (Chao et al., 2010; Durand et al., 2012; Han et al., 2012; Hensch, 2005; Hensch et al., 1998; Huang et al., 1999; Krishnan et al., 2015; Lupori et al., 2019; Mazziotti et al., 2017; Picard and Fagiolini, 2019; Pizzo et al., 2016; Terzic et al., 2021; Wen et al., 2018). Here, we present the first systematic characterization of PNN expression in the male mouse model of CDKL5 deficiency disorder with subregion-, anatomical-, and hemisphere-specificity during visual cortex critical period (**Fig. 8**). Our findings within the *Cdkl5*-null male primary visual cortex agree with data from the V1B representative section analyses performed in Pizzo et al. 2016. Extended analysis in V1B, V1M and rostral V1 over critical period also point to the earliest changes in dysregulated WFA expression at P15, indicating precocious atypical PNN expression in the *Cdkl5*-null male visual cortex (**Fig. 8A**). Particularly, the consistent and early changes in the V1M and V1B are novel results, with implications for the imbalance between excitation and inhibition in the *Cdkl5*-null visual cortex. This finding aligns well with previously reported precocious critical period closure in the *Mecp2*-null visual cortex, which led to immature binocular matching patterns (Krishnan et al., 2015), and depth perception issues (Durand et al., 2012). Such atypical inhibitory changes to the neural network are likely to contribute to the visual impairments associated with CDKL5-deficiency disorder pathology. Further electrophysiological studies are necessary to determine how subregion-specific PNN expression differences (**Fig. 3, 7**) ultimately contribute to the differentially evoked activity, *in vivo* neural activity and/or synchrony in the *Cdkl5*-null visual cortex across development.

Subregional differences are particularly important in the context of the rostral-caudal positioning, with P30 null V1M showing fewer caudal PNNs, while rostral V1B of P30 null males display increased high-intensity PNNs (vs P30 WT; **Fig. 4**). Such positional information points to nuanced and likely compensatory synaptic plasticity mechanisms in the *Cdkl5*-null brain. Additionally, as parvalbumin+ interneuron networks influence PNN maintenance, it is likely that this network activity and/or gamma oscillations are disrupted in a subregion-specific manner in the *Cdkl5*-null visual cortex (Bartos et al., 2002; Devienne et al., 2021). Future work should determine the impact of classical critical period manipulations (e.g.: monocular deprivation, dark rearing) on PNN expression to determine the role of CDKL5 protein during this sensitive period. Conditional knockout of *Cdkl5* in glutamatergic neurons produces robust neurological phenotypes, indicative of a more significant role for CDKL5 in select neuronal subtypes over others (Awad et al., 2023; Lupori et al., 2019; Schroeder et al., 2019). From a translational or therapeutic perspective, exciting new biomarkers and candidates have been proposed in recent works (Saby et al., 2022; Van Bergen et al., 2022). However, as most of the above studies focus predominantly on null male mice (*including the present study*), further in-depth analyses across age are required using the female *Cdkl5*-heterozygous mouse model (Fuchs et al., 2018). Due to the subtle PNN phenotypes detected in null male mice, we could not justify performing similar visual cortical analysis in the *Cdkl5*-heterozygous female mice, due to the inherent mosaicism and lack of reliable antibodies to label CDKL5 protein in individual cells of the heterozygous females (unpublished data from the lab).

Previously, we reported that the adult female primary somatosensory cortex exhibited lateral-medial axis and hemisphere-specific changes in high-intensity PNN density, in both WT and *Mecp2*-heterozygous mice (Lau et al., 2020b). Such nuanced discovery was possible due to systematic whole brain analysis. We speculated that such anatomical differences could contribute to, or be shaped by, the functional specialization of the cortex. This functional specialization in the barrel cortex could contribute to the individual mouse’s whisker side preference, similar to handedness in primates (Lonsdorf and Hopkins, 2005). Determining when and how PNN lateralization occurs and is maintained across cortical subregions will require much work and seed new directions for the field. Toward this goal, we recently identified that adolescent 6-week-old female WT and *Mecp2*-heterozygous do not exhibit hemisphere laterality in PNN expression within the primary somatosensory cortex (Mykins et al., 2023), suggesting asymmetry in PNN density in the adult primary somatosensory cortex is an age-related, and likely experience-dependent, cellular phenotype. During these adolescent and early adulthood phases, it is likely that tactile sensory perception is undergoing a sensitive period, influenced by puberty and gonadal hormones. Thus, it is likely that the maturation of PNN density and lateralization in expression in this cortical subregion is not finalized yet. Since the visual critical period matures earlier, we hypothesized an earlier lateralization of PNN expression in the primary visual cortex, though the functional relevance of such lateralization in rodents is unclear. Humans, on the other hand, are known to have a “dominant” eye (Chan and Chang, 2022; Dieter et al., 2017; Ooi et al., 2013; Ooi and He, 2020; Yang et al., 2010). However, we did not observe hemisphere or anatomical axes specific laterality in PNN density in the mouse primary visual cortex at P15 or P30. Additionally, this lack of lateralization was preserved in the *Cdkl5*-null male mice, indicating that CDKL5 is not required for establishing or maintaining this lateralization in developing male mice.

## CONFLICT OF INTERESTS

The authors declared no conflict of interests.

## AUTHOR CONTRIBUTIONS

KK supervised the project, designed experiments and analyzed data. BE and ME perfused animals, performed sectioning, immunostained, mapped sections, and manually counted PNNs. BE imaged sections. LRD performed PNN histogram analysis. ME, LRD and BYBL performed data analysis and data visualizations. BYBL trained BE and ME in technical aspects of the project. BE, ME, LRD, BYBL and KK wrote and edited the manuscript.

## ACKNOWLEDGEMENTS

We would like to thank the following undergraduate students for their help with PNN counting and other technical contributions: Sarah-Anne Bowyer and Bailey Bohannon. This work was supported by National Science Foundation Graduate Research Fellowship Program (GRFP) (LRD), by research assistantships to BE and ME from the University of Tennessee, Knoxville, through startup funds to KK and by the National Institute of Mental Health of the National Institutes of Health under Award Number R15MH124042 (KK).

## Notes

### Competing Interest Statement

The authors have declared no competing interest.

### Summary of Updates

We performed new histogram analyses of PNN expression and added another developmental time point, postnatal day 15. These results culminated to two new graphs and a revised summary diagram. We also added another author, Logan Reid Dunn.

